# Temporal consistency of judgement biases in bumblebees

**DOI:** 10.64898/2026.04.07.716885

**Authors:** Luigi Baciadonna, Vivek Nityananda

## Abstract

Judgment bias tasks are increasingly used to assess affective states in animals, yet the extent to which they might reflect transient states or stable traits remains unclear. Here, we tested bumblebees (*Bombus terrestris*) in an active choice task across three repeated sessions to assess individual consistency in the absence of any manipulation. Bees were trained to associate each of two colours with either a high or a low reward, presented in separate chambers. During testing, they were presented with ambiguous colours. Bees were more likely to choose the high reward chamber and to choose more quickly in response to colours closer to the positive colour. The latency to choose the cues showed significant and moderate repeatability across sessions, suggesting a stable, trait-like underlying component. In contrast, the repeatability of the chamber choices was negligible, indicating that such responses might be largely state-dependent and influenced by situational factors. These findings suggest that judgment biases, particularly as assessed through an active choice task reflect states affected by external factors. Active choice tasks may help disentangle stable behavioural traits from transient affective states in invertebrates.

## Introduction

Reliable measures for assessing animal emotions are essential, especially when they can inform and guide welfare-related decisions or contribute to translational research. A well-known paradigm to assess these states is the judgement bias test, where emotional states are inferred based on the evaluation of an ambiguous stimulus [1,2]. In humans, studies employing this paradigm have demonstrated that people with depression or anxiety disorders tend to interpret ambiguous situations negatively [3,4]. Over the last decades, this paradigm has been adapted to non-human animal species, including invertebrates [5,6].

Since the seminal study [7] that tested this approach in rats, several studies have applied the same approach across multiple species. Research on invertebrates, specifically honeybees, bumblebees, and fruit flies, demonstrates that they are malleable to the experimental induction of positive or negative emotional states, as demonstrated by cognitive biases in response to ambiguity [6]. A recent meta-analysis of 71 studies across 22 species supports the overall validity of experimental manipulations to probe affective states in animals [5]. However, the effect sizes are relatively small and appear to be affected by several methodological and biological moderators, including task type, training cue reinforcement, and the sex of the animals.

Task choice, for example, can impact findings and their interpretation, as suggested by differences between studies using a go/no-go or active choice paradigm [5,8,9]. Go/no-go paradigms require responding to an ambiguous stimulus after learning to respond to a positive one and suppressing the response to a negative one. Active choice paradigms rely on active responses for each stimulus. Since the animal must make a clear choice, active choice paradigms mitigate against the influence of confounding factors, specifically motivational changes, thus increasing the validity and interpretability of the results [5].

While judgment bias tasks are widely used to assess animal affect, the responses may reflect either transient states or stable traits [5,9,10]. Contextual factors, including housing, stressors, or mood-altering drugs can modulate the bias, indicating situational changes typical of an emotional change [5,11]. At the same time, individual animals can display consistent predispositions to interpret ambiguous stimuli optimistically or pessimistically, suggesting a trait component [11]. Similar patterns have been reported in pigs, companion animals, and wild species [16–22]. Baseline differences across breeds, test designs, or motivational contexts further complicate interpretation [23–26]. Assessing repeatability can help separate these aspects. While state and trait effect are not mutually exclusive, quantifying repeatability provide a baseline estimate of behavioural variability and provides an estimate of how much variation is due to consistent individual differences. This baseline is crucial for meaningful interpretation. Even when treatment effects are detected, their magnitude and reliability may depend on the level of underlying variability. Despite this, repeatability of judgement biases has been investigated only in a few species, showing that individual consistency can range from stable to highly variable [13,27–29]. Even studies without manipulations are rare; two go/no-go studies found strong learning effects across sessions [27,30]. Thus, the extent to which judgment biases reflect transient affective states rather than stable traits remains unclear.

We used an active choice judgment bias test to measure the effect of repeated testing of judgement bias in bumblebees (*Bombus terrestris*). We hypothesised that, in the absence of any manipulation, responses to ambiguous stimuli would remain stable across repeated tests, indicating a trait component of judgment bias in bumblebees.

## Material and Methods

### Animals and experimental set-up

The study was conducted on female worker bumblebees (*Bombus terrestris*). To control for age, only individuals tagged within 24 h of eclosion were included. When not tested, bees had ad libitum access to a 20% (w/w) sucrose solution and approximately 3 g of pollen per day. During experiments, bees were presented with solid-colour visual stimuli on a monitor in the flight arena (***figure 1a***), using the colours detailed in Procenko et al. [12]. Studies on Hymenoptera are not regulated under the Animals (Scientific Procedures) Act 1986 (ASPA). See supplementary materials for additional details.

**Figure 1.**
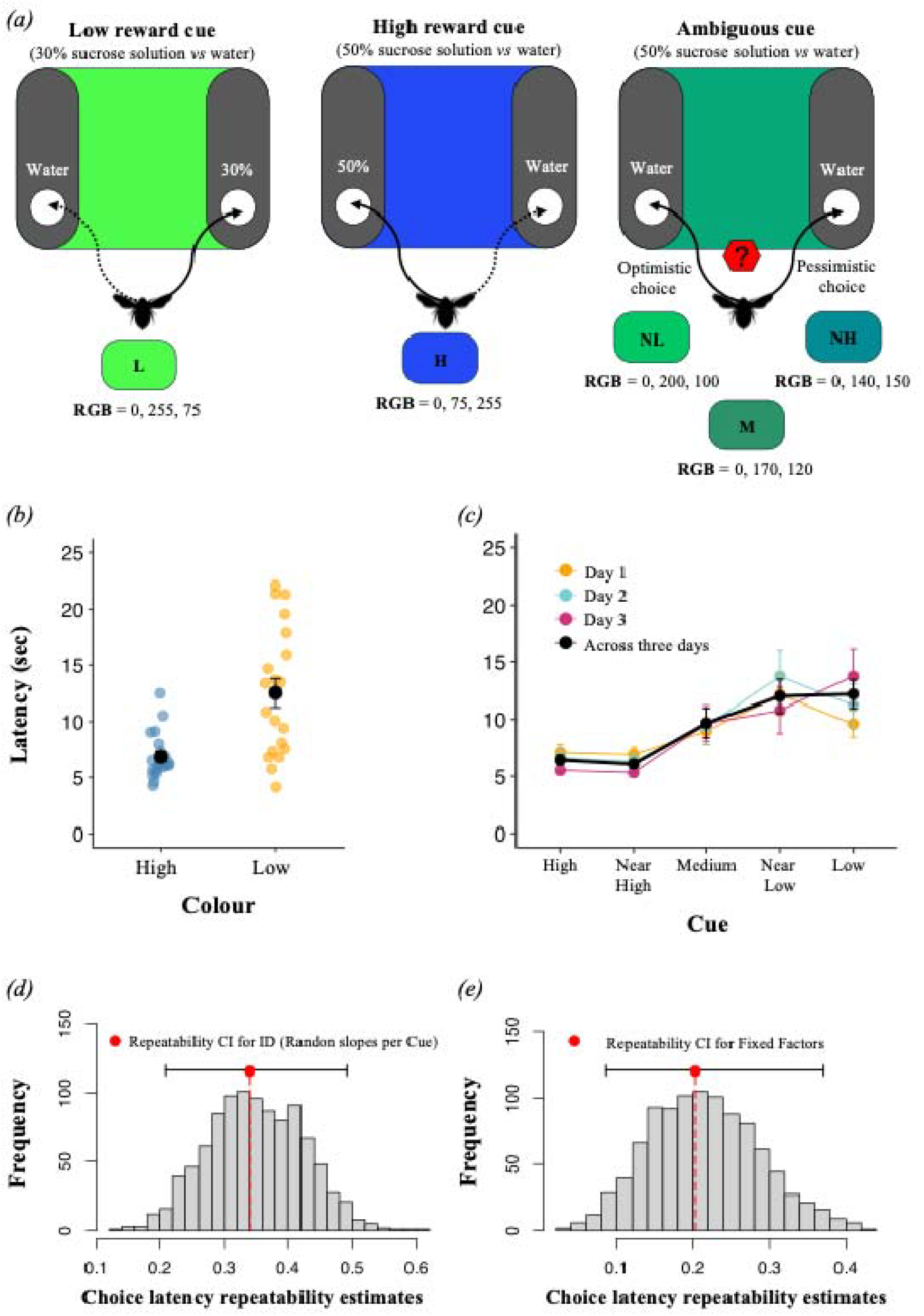
Experimental paradigm and choice latency analyses. ***(a)*** An example scenario during bee training, with green indicating a low reward (a 30% sucrose solution) in the right chamber, and blue indicating a high reward (a 50% sucrose solution) in the left chamber. In both cases, the other chamber would contain only distilled water. The association between colour, reward and location was counterbalanced across trials. The test phase consisted of five trials in which different colours were presented on the screen in a pseudorandom order (cue values: L, low; NL, near low; M, medium; NH, near high; H, high). ***(b)***Latency to approach high and low rewards at the end of training. ***(c)*** Response latency to each of the five cues during tests. Each colour represents a different day and the black line is the average across all days. The error bars represent the standard error of the mean. ***(d and e)*** Histograms of repeatability estimates (R) of choice latency for (*d*) bee identity and (*e*) fixed factor. The histograms represent the distribution of R values generated from 1.000 bootstrap iterations. The R value represents the proportion of total variance explained by individual identity (ID) in (*d*) and by the fixed effect included in the model in (*e*). In both plots, the dot and dashed line represent the point estimate. For bee identity (*d*), the estimate (R = 0.34) indicates that approximately 34% of the variation in speed was due to consistent differences between individual bees. Similarly, the estimate for fixed effects (R = 0.20) indicates that approximately 20% of the variance in choice latency was attributable to the fixed factor included in the model.

### Pretraining

Before conditioning, bees were familiarised with the reward chambers and allowed to perform independent foraging bouts. Only bees that reliably located the reward without assistance proceeded to training. See supplementary materials for additional details.

### Training

The conditioning procedure was identical to the one described previously [12]. Two colours (blue and green) were each associated with a different sucrose concentration (low: 30% w/w; high: 50% w/w) in one of two reward chambers (***figure 1a***). On each trial one of two colours was presented. One colour indicated that a reward (either high or low value) was available in the chamber on one side of the apparatus, while the opposite chamber contained distilled water. For the other colour, the reward and the water location were reversed. Colour, reward value and side were counterbalanced across bees (table S1). Training continued until individuals reached ≥80% correct choices in the last 20 trials. A total of 24 out of 25 bees passed the initial conditioning, with one bee excluded due to strong side biases. Only full entries, when the bee’s body fully entered the chamber, were recorded. All trials were video recorded and choice latencies were calculated as the mean of the last two training trials for each respective colour. See supplementary materials for additional details.

### Judgement bias testing

The test phase consisted of five trials, each presenting a different colour cue on the monitor. Test colours included the two conditioned colours (green and blue) and three ambiguous colours of intermediate value: near-blue (RGB: 0, 140, 150), medium (RGB: 0, 170, 120), and near-green (RGB: 0, 200, 100). Depending on their proximity to the colour indicating a high- or low-reward, ambiguous cues were classified as near-high, medium, or near-low (***figure 1a***). The order of colour presentation was pseudorandomized across bees, with the first test colour always being one of the three ambiguous colours. During testing, all cues were unrewarded, with both chambers containing 0.2 ml of distilled water. After the bee made its first choice, it was gently captured and returned to the tunnel connecting the nest box and the arena. Between presentations of test cues, bees received refresher trials consisting of two presentations of each conditioned colour with the corresponding reward. If a bee made an incorrect choice, an additional set of four trials was administered. This procedure was repeated up to two more times (for a total of three sets of four trials), and if errors persisted, refresher training continued until the bee met the learning criterion (≥ 80% correct choices in the last 20 trials). All trials were video recorded. Bees’ first choices, defined as fully entering a reward chamber, and the choice latency, measured as the time from leaving the tunnel to entering the reward chamber, were recorded.

### Judgement bias testing across consecutive days

The judgement bias test was carried out on the following two days as described above. Each session began with four refresher trials (two presentations of each conditioned colour with its reward). If a bee made an incorrect choice, the same steps described above were followed until the bee reached the learning criteria. After the refresher trials, tests were carried out as above. From the initial pool of 24 bees, four did not complete all tests because they died. Specifically, one bee completed only the first test and three bees completed two out of the three planned tests.

### Statistical analyses

Our hypothesis and statistical analyses were preregistered at aspredicted.org (220930). The data were plotted and analysed using RStudio v. 3.2.2 (R Foundation for Statistical Computing, Vienna, Austria). Subsequent statistical models were fitted by maximum-likelihood estimation. Models were compared using the “model.sel” function in the MuMIn package [13] and the model with the lowest Akaike information criterion (AIC) score was selected as the best model. We used the package DHARMa [14] for residual testing of all models. The p-value of each factor was derived using the ‘‘drop1’’ function [15] and post-hoc analysis were performed using “emmeans” function [16].

Latency data in all analyses were log-transformed and latencies greater than 1.5 times the interquartile range were excluded (18 out of 120 datapoints for training, 46 out of 300 for tests). Choice latency for the last two training bouts was analysed using a linear mixed-effects model (lmer function, lme4 package), including only bees that completed all three tests [17]. The model included “Colour” (categorical: High or Low), “Training Day” (categorical: 1, 2 and 3) and their interaction, “Age”, “High Colour” (categorical: Blue or Green), “High side” (categorical: Right or Left, position of the high reward) and “Colony” (categorical: 1, 2 and 3) as fixed factors. Bee identity (‘ID’) was included as a random intercept variable.

The model analysing choice latency for the judgement bias test included “Cue” (categorical: High, Low, Near High, Medium and Near Low), Testing Day” (categorical: 1, 2 and 3) and their interaction, “Age”, “High Colour Side” (categorical: Blue/Right, Blue/Left, Green/Right and Green/Left) and “Colony” (categorical: 1, 2 and 3) as fixed factors. Bee identity (“ID”) was included as a random intercept variable.

To assess the consistency of individual differences in latency to choose across days and cues, we estimated repeatability using the rptR package [18]. The model included the same fixed factor with individual identity (“ID”) specified as a random effect with random slopes for “Cue”. Repeatability was calculated as the proportion of total variance in choice latency explained by differences between individuals. Confidence intervals were obtained using 1000 parametric bootstrap iterations. The variance explained by fixed effects was also estimated to quantify their contribution relative to individual consistency.

Choices of reward chambers previously associated with high and low value cues were coded as 1 or 0 respectively. The choices of high-reward chambers were classified as optimistic, and choices of low-reward chambers as pessimistic. We analysed the probability of optimistic choice with a generalized linear mixed-effect model (glmer function, lme4 package) with binomial distribution and a logit link function. The fixed factors were ‘Cue” (categorical: High, Low, Near High, Medium and Near Low), “Testing Day” (categorical: 1, 2 and 3) and their interaction, “Age”, Order (categorical: 1, 2, 3, 4, 5 representing the sequence of colour cue presentations) and “Colony” (categorical: 1, 2 and 3). Bee identity (‘ID’) was included as a random intercept variable. Repeatability of individual choices was estimated using the rptR package [18]. Due to lack of convergence using the model described above, repeatability was assessed using a model which included only the random intercept for bee identity and the binary response variable. Repeatability was calculated as the proportion of total variance explained by differences between individuals, and confidence intervals were obtained via 1000 bootstrap iterations. This analysis provided an estimate of how consistent bees were in their optimistic choices across trials.

## Results

### Training

During training, bumblebees achieved the learning criterion and continued to the judgment bias test. The best model (AIC = 134.6) for choice latency data included the main effects of colour cue (GLMM, χ ^2^ (1) = 37.20, p = 3.386^e-08^). Bees were significantly faster to choose a high reward colour compared with a low reward colour in the last training choice (estimate ± s.e. = - 0.50 ± 0.08, t = - 6.05, p < 0.0001; ***figure 1b***). The difference in latencies indicated that the bees could differentiate between both the colour cues and the rewards. Including the full dataset, which contains latencies exceeding 1.5 times the interquartile range, produced similar results (table S2).

### Judgement bias tests

The best model (AIC = 346.1) for the test choice latency data included the main effect of cue. Cue significantly affected bee latency to choose a chamber (GLMM, χ ^2^(4) = 22.89, p = 5.497^e-16^; ***figure 1c***), with bees showing progressively longer latencies for colours closer to that associated with lower reward (High versus Medium: estimate ± s.e. = - 0.33 ± 0.08, t = - 4.00, p = 0.0008; High versus Near Low: - 0.55 ± 0.08, t = - 6.54, p < 0.0001; High versus Low: - 0.56 ± 0.08, t = - 6.35, p < 0.0001; see supplementary materials for additional details). Including the full dataset, which includes latencies exceeding 1.5 times the interquartile range, showed similar results (table S3). Repeatability analysis indicated that individual bees were moderately and significantly consistent in their latency to choose (R = 0.34 ± 0.07, 95% CI = 0.20 – 0.49, p = 1.23^e-07^; ***figure 1d***), reflecting stable individual differences (see supplementary material for results including the full dataset). Fixed effects explained a comparable proportion of variance (R = 0.20 ± 0.07, 95% CI = 0.08 – 0.37; ***figure 1e***). Bees’ latencies to choose across testing days for each cue are plotted in ***figure 2a***.

**Figure 2.**
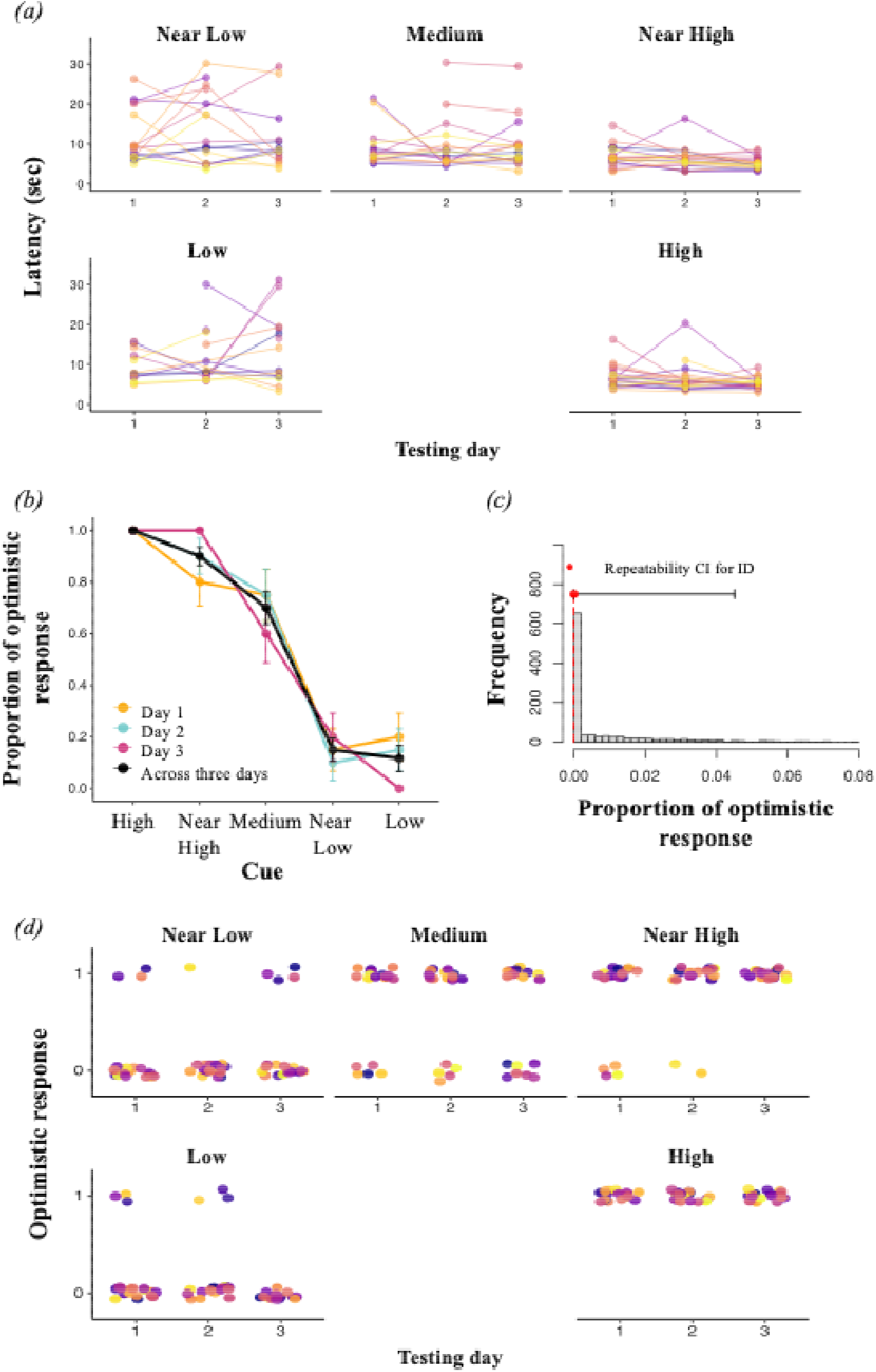
Bee choice analyses. ***(a)*** Individual bee choice latencies in tests across testing days for each cue. Each colour and line depicts a different individual. ***(b)*** The proportion of bees that made an optimistic choice in response to each of the five cues. Other details as in Fig 1d.***(c)*** Histogram of repeatability estimates of the optimistic response ***(d)*** Individual bee optimistic responses across testing days for each cue.

The best model (AIC = 214.7) for reward chamber choice included the main effect of cue. Cue significantly influenced the likelihood of bees making an optimistic choice (GLMM, χ^2^ (4) = 206.72, p = 2.2^e-16^; ***figure 2b***). Bees were progressively less likely to respond optimistically to cues further from the High cue. They made more optimistic responses to the Near High cue than the Near Low (estimate ± s.e. = 4.30 ± 0.59, z = 7.20, p < 0.0001) and Low cues (4.61 ± 0.62, z = 7.35, p < 0.0001; see supplementary materials for additional details). Bees always responded optimistically to the High cue. Repeatability analysis across all cues indicated no detectable variance attributable to individual identity (R = 0, SE = 0.013, 95% CI = 0–0.044, p = 1; ***figure 2c***). We therefore excluded the conditioned cues, leaving only the ambiguous ones (Near High, Medium, Low). However, the repeatability remained negligible (R = 0, SE = 0.022, 95% CI = 0–0.078, p = 0.5). These results indicate that bees responded similarly to each cue, leaving little inter-individual variation for the model to detect (***figure 2d***). To further support this, we calculated the coefficient of variation (CV) for responses to each cue, which showed that bees’ responses were highly uniform for the conditioned extreme cues (High: mean ± SD; CV = 1 ± 0, CV = 0; Low: 0.10 ± 0.31, CV = 2.94;) and moderately variable for ambiguous cues (Near High: 0.89 ± 0.31. CV = 0.34; Medium: 0.70 ± 0.46, CV = 0.65; Near Low: 0.14 ± 0.35, CV = 2.50).

## Discussion

We tested the hypothesis that a judgment bias reflects a stable individual trait rather than a transient affective state and that responses to cues would therefore remain consistent across multiple tests. Our repeatability analyses of behaviour in judgement bias tests revealed moderate consistency in individual latency to choose but negligible repeatability in the probability of an optimistic choice in response to ambiguous stimuli.

An important aspect in judgment bias paradigms, especially in go/no-go tasks, is that subjects often learn that ambiguous cues are not rewarded, which can progressively reduce their ambiguity and lead to increased latency and number of no responses across repeated sessions [9,19–21]. Secondary or partial reinforcement have been suggested to mitigate the loss of ambiguity [22–26]. However, we argue that the use of active choice tests is a convenient and effective method to avoid learning effects and other confounding factors. We found no interaction between cue and testing time for either latency or optimistic choice, indicating that bees’ responses to learned and ambiguous cues remained stable over time. This suggests that repeated exposure did not lead bees to learn that the ambiguous cues were unrewarded, maintaining their uncertainty across days. The persistence of graded responses further supports the idea that the active choice task, along with the refresher trials performed during the test, were effective in preserving cue ambiguity. These results demonstrate the potential benefit of using an active choice task for assessing judgment bias over repeated tests, as it reduces the confounding effects of learning and motivation often reported in go/no-go tasks [12].

In our study, bees showed moderate repeatability in their latency to choose across cues (R = 0.34), with some individuals being consistently faster or slower across repetitions. Such repeatability aligns with the level of consistency found in literature, where typically 37% of individual variation is attributed to stable individual differences [27]. This suggests that choice latency may underpin a stable trait supporting the covariation hypothesis between cognition and personality [28–31]. Differences in personality traits may underpin emotional states, and this relationship can be bidirectional. Individuals with a specific personality type can be more prone to experience positive or negative affective states, leading to corresponding cognitive biases. At the same time, repeated exposure to negative or positive experiences may change these traits. The observed stability in choice latency suggests that while decision speed may be driven by stable metabolic and physiological constraints, optimistic choice i.e. judgment remains flexible [32,33,33]. This distinction is ecologically important as it allows bees to maintain consistent activity levels while modulating their evaluation of ambiguous cues in response to fluctuating rewards. To further validate the judgment bias test, it would be valuable to include additional behavioural measures that are hypothesised to covary with emotional states.

Repeatability of optimistic choices for ambiguous cues was negligible, suggesting that the likelihood of making an optimistic choice fluctuated across the three repetitions, whereas responses to the trained cues, based on the coefficient of variation, were highly consistent. In contrast to latency, optimistic responses appear to be largely state-dependent and influenced by situational factors rather than stable individual traits. The absence of consistent individual biases indicates that bees were unbiased in the absence of any experimental manipulation. Therefore, when a manipulation elicits a detectable shift in active choices in a judgment bias test, such a change is more likely to reflect state-dependent variation rather than a stable individual difference, supporting an interpretation of them as a shift in affective states. For instance, we found that the variability in the likelihood of making an optimistic choice across three testing days was lower than the variation observed when bumble bees were shaken or trapped in a comparable experimental set-up [11]. This suggests that variability in the absence of manipulation is substantially lower than when the animals’ affective state is experimentally altered.

In conclusion, our findings demonstrate that repeated testing can be performed without confounding effects of learning and support the use of active choice tasks to assess transient and stable emotional states in invertebrates.

## Acknowledgements

LB and this research are supported by a Leverhulme Trust Research Project Grant (RPG-2021-358).

## Notes

### Competing Interest Statement

The authors have declared no competing interest.

