## Supplementary Materials for "Temporal consistency of judgement biases in bumblebees"

**Temporal consistency of judgement bias in bumblebees: Supplementary Materials**

**This file includes:**

Supplementary Methods and Results

**Methods**

*Animals and experimental set-up*

Bumblebees were obtained from a commercial supplier (Agralan, UK). Upon arrival in the laboratory, each bee was individually tagged for identification using numbered colour tags (Opalith Zeichenplättchen Leuchtfarben, Bienen-Voigt & Warnholz, Ellerau, Germany). Tagging involved gently restraining each bee in a marking cage, pressing it against the mesh with a sponge, and glueing the tag to the dorsal thorax (Loctite Super Glue Power Gel). Marked bees were then transferred to an artificial two-compartment plastic nest box (28 × 16 × 12 cm). One compartment housed the colony, while the other contained cat litter to function as a waste area. The nest box was connected to a flight arena (110 × 61 × 40 cm) via a transparent acrylic tunnel (56 × 5 × 5 cm). The arena was illuminated through a UV-transparent Plexiglas® lid (Philips HF-P 114-35 TL5 ballast, daylight fluorescent tubes, Osram, Germany). Although not regulated under Animal Act 1986 all housing and experimental procedures nonetheless followed the 3Rs principles (see <http://www.nc3rs.org.uk/>), and the study was not invasive.

Visual stimuli consisted of full-screen, solid-colour displays presented on a 24-inch LED monitor (Dell U2412M, 1920 × 1200 px). Stimulus presentation was controlled via custom MATLAB scripts (MathWorks, Natick, MA, USA) using the PsychToolbox package [1]. The colour of the visual stimuli selected and validated were the same used in Procenko et al. [2]. Two vertical panels (40 × 8 cm) were positioned 8.5 cm in front of the left and right edges of the monitor, leaving the centre visible. Each panel contained an opening that held a glass vial (7 ml, 10 mm internal diameter) serving as a reward chamber, positioned 7 cm above the arena floor. After each visit, the reward chambers were substituted with a new one to eliminate any residual scent marks or pheromonal cues. At the end of each experimental day, all equipment was washed in hot water followed by 70% ethanol and air-dried before reuse.

*Pretraining*

Prior to training, individual bees were familiarised with both reward chambers. Each bee was captured at the exit of the tunnel connecting the nest box to the flight arena using a plastic cup, which was then aligned to the entrance of one of the two reward chambers containing a droplet of sucrose solution (0.2 ml, 30% w/w). Bees were then allowed to return to the nest box and on subsequent visits, they could find the reward chambers independently. Bees that reliably located the reward and performed repeated foraging bouts without assistance were selected to start training.

*Training*

Bees were presented one of two colours on an LED screen, green (RGB: 0, 255, 75) or blue (RGB: 0, 75, 225), each associated with a different sucrose concentration (low: 30% w/w; high: 50% w/w) in one of two reward chambers. On each trial, only one colour was presented, indicating either a high or low value reward in the chamber on one side, with distilled water in the unrewarded chamber. When the other colour was presented, the other reward (low or high) was presented in the chamber on the opposite side, again with distilled water in the unrewarded chamber. For instance, a bee could be conditioned to green indicating high reward on the left chamber and blue indicating low reward on the right chamber. Bees experienced the rewarded and unrewarded colours an equal number of times, and the order of trials was pseudorandomized so that no colour–reward combination occurred more than twice in a row. The combination of colour, side and reward were counterbalanced across all bees. Training continued until bees reached the learning criterion of ≥80% correct choices in the last 20 trials. All trials were video recorded from above using a mobile phone camera (Huawei Nexus 6P, 1440 × 2560 px, 25 fps).

**Table S1. Colour, reward value, chamber side and bees excluded across the sample size.**

| **Colour – Reward value – Side** | **N of bees** | **Excluded** |
| --- | --- | --- |
| Blue High Left / Green Low Right | 6 | 1 |
| Blue High Right / Green Low Left | 5 |  |
| Green High Left / Blue Low Right | 7 | 2 |
| Green High Right / Blue Low Left | 6 | 1 |

**Results**

*Judgement bias tests*

Pairwise comparisons showed that choice latencies for the Near High cue differed significantly from those for the Medium (estimate ± s.e. = - 0.36 ± 0.08, t = - 4.41, p = 0.0002), Near Low (- 0.59 ± 0.08, t = - 6.91, p < 0.0001), and Low cues (- 0.60 ± 0.08, t = - 6.71, p < 0.0001). No significant differences were detected among the remaining cue pairs (p ≥ 0.07 in all cases).

Pairwise comparisons also showed that Medium cue elicited more optimistic responses than the Near Low (2.86 ± 0.48, z = 5.92, p < 0.0001) and Low cues (3.17 ± 0.52, z = 6.10, p < 0.0001). No significant differences were detected among the remaining cue pairs (p ≥ 0.057 in all cases).

The repeatability analysis indicated that individual bees were also moderately and significantly consistent in their latency to choose even when the full model includes latencies exceeding 1.5 times the interquartile range (R = 0.29 ± 0.06, 95% CI = 0.16 – 0.43, p = 1.63^e-10^), reflecting stable individual differences. Fixed effects explained a comparable proportion of variance as in the model excluding outliers (R = 0.23 ± 0.05, 95% CI = 0.12 – 0.35).

**Table S2. Summary of the generalized linear mixed model examining the influence of the fixed factors on choice latency using the full database (including the latencies greater than 1.5 times the interquartile range) for the training data.** The best model (AIC 240.2) for choice latency included only the main effects of colour (GLMM, χ ^2^_(1)_ = 56.33, p = 3.25^e-11^). The *post-hoc* revealed that bees are significantly faster to choose a high reward colour compared to a low reward colour in the last training choice (estimate ± s.e. = - 0.87 ± 0.11, t = - 7.46, p < 0.0001).

| Response variable: Log(Latency) | | | | | |
| --- | --- | --- | --- | --- | --- |
| Model: Log(Latency) ~ Colour + (1 \| ID) | | | | | |
| **Fixed Effect** | **Estimate** | **SE** | **df** | **t value** | **P value** |
| Intercept (High) | 1.85 | 0.089 | 51.27 | 20.86 | < 2^e-16^ |
| Colour (low vs High) | 0.87 | 0.116 | 95.04 | 7.51 | 3.26^e-11^ |

**Table S3. Summary of the generalized linear mixed model examining the influence of the fixed factors on choice latency using the full database (including the latencies greater than 1.5 times the interquartile range) for the test data.** The best model (AIC 733.8) for choice latency included only the main effects of cue (GLMM, χ ^2^_(4)_ = 29.65, p = 2.2^e-16^).

| **Fixed Effect** | **Estimate** | **SE** | **t value** | **P value** |
| --- | --- | --- | --- | --- |
| Intercept | 1.77 | 0.13 | 13.43 | < 2^e-16^ |
| Cue Near High | 0.03 | 0.14 | 0.25 | 0.79 |
| Cue Medium | 0.61 | 0.14 | 4.40 | 1.52^e-05^ |
| Cue Near Low | 0.92 | 0.14 | 6.58 | 2.24^e-10^ |
| Cue Low | 1.22 | 0.14 | 8.73 | 2.45^e-16^ |

**Table S4. Summary of post-hoc analysis for fixed factors Cue for the full database (including the latencies greater than 1.5 times the interquartile range) for the test data.** The *post-hoc* tests revealed that bees are significantly faster to choose a high reward colour compared to a low reward colour in the last training choice (estimate ± s.e. = - 0.87 ± 0.11, t = - 7.46, p < 0.0001).

| **Contrast** | **Estimate** | **SE** | **T value** | **P value** |
| --- | --- | --- | --- | --- |
| High vs Near High | - 0.03 | 0.14 | - 0.25 | 0.99 |
| High vs Medium | - 0.61 | 0.14 | - 4.37 | 0.0002 |
| High vs Near Low | - 0.92 | 0.14 | - 6.54 | < 0.0001 |
| High vs Low | - 1.22 | 0.14 | - 8.67 | < 0.0001 |
| Near High vs Medium | - 0.58 | 0.14 | - 4.12 | 0.0005 |
| Near High vs Near Low | - 0.88 | 0.14 | - 6.28 | < 0.0001 |
| Near High vs Low | - 1.18 | 0.14 | - 8.41 | < 0.0001 |
| Medium vs Near Low | - 0.30 | 0.14 | - 2.16 | 0.19 |
| Medium vs Low | - 0.60 | 0.14 | - 4.29 | 0.0002 |
| Near Low vs Low | - 0.30 | 0.14 | - 2.12 | 0.21 |
